# Neuronal dynamics of the default mode network and anterior insular cortex: Intrinsic properties and modulation by salient stimuli

**DOI:** 10.1101/2022.08.01.501899

**Authors:** Tzu-Hao Harry Chao, Byeongwook Lee, Li-Ming Hsu, Domenic Hayden Cerri, Wei-Ting Zhang, Tzu-Wen Winnie Wang, Srikanth Ryali, Vinod Menon, Yen-Yu Ian Shih

**Author notes:** These authors contributed equally to this work (Tzu-Hao Harry Chao, Byeongwook Lee, Li-Ming Hsu). Equal contribution (Vinod Menon, Yen-Yu Ian Shih). Correspondence to: Yen-Yu Ian Shih, Ph.D. and Vinod Menon, Ph.D.

## Abstract

The default mode network (DMN) is closely associated with self-referential mental functions and its dysfunction is implicated in many neuropsychiatric disorders. However, the neurophysiological properties and task-based functional organization of the rodent DMN are poorly understood, limiting its translational utility. Here, we combine fiber-photometry with fMRI and computational modeling to characterize dynamics of putative rodent DMN nodes and their interactions with the anterior insular cortex (AI) of the salience network. We reveal neuronal activity changes in AI and DMN nodes prior to fMRI-derived DMN activations and uncover cyclical transition patterns between spatiotemporal neuronal activity states. Finally, we demonstrate that salient oddball stimuli suppress the DMN and enhance AI neuronal activity, and that the AI causally inhibits the retrosplenial cortex, a prominent DMN node. These findings elucidate previously unknown properties regarding the neurobiological foundations of the rodent DMN and its modulation by salient stimuli, paving the way for future translational studies.

**Highlights:**

- Concurrent measurement of neuronal (GCaMP) and fMRI signals in retrosplenial, cingulate, prelimbic, and anterior insula cortices
- GCaMP signals reveal neuronal antagonism between AI and fMRI-derived DMN activation and deactivation
- GCaMP signals reveal salient oddball stimuli-induced suppression of prelimbic, cingulate and retrosplenial cortices, and activation of anterior insular cortex
- Anterior insular cortex causally inhibits retrosplenial cortex during processing of salient oddball stimuli
- Findings delineate neurofunctional organization of the rodent DMN and provide a more informed model for translational studies

## Introduction

Discovery of the default mode network (DMN) in 2003 (Greicius et al., 2003) sparked significant interest in the large-scale functional organization of the human brain (Raichle, 2015). The DMN comprises brain areas that are consistently deactivated during a wide range of cognitively demanding tasks, notably these regions also demonstrate highly synchronous activity during task-negative “resting-state” conditions (Anticevic et al., 2012; Greicius and Menon, 2004; Greicius et al., 2003; Menon, 2011; Seeley et al., 2007; Shulman et al., 1997). Seminal fMRI studies in humans have identified the retrosplenial cortex (RSC), posterior and rostral anterior parts of cingulate cortex (Cg), medial prefrontal cortex (mPFC), and the inferior parietal lobe (IPL) as key nodes of the DMN (Buckner et al., 2008; Chase et al., 2020; Eickhoff et al., 2016; Greicius et al., 2003; Raichle, 2015; Reid et al., 2016; Sridharan et al., 2008). In humans, the DMN is thought to play a fundamental role in self-referential mental functions, including recollection of autobiographical events (Buckner et al., 2008) and understanding the mental states of others (Garrison et al., 2015; Molnar-Szakacs and Uddin, 2013). Dynamic changes in DMN activation have also been associated with the mediation of perception and cognition (Das and Menon, 2020; Menon, 2011, 2013; Sala-Llonch et al., 2012; Schaefer et al., 2014; Smith et al., 2018). While brain imaging studies have provided significant insights into the human and rodent DMN (Lang et al., 2014; Menon, 2011, 2013; Smitha et al., 2017), fMRI does not directly measure neuronal activity, consequently little is known about the neuronal processes underlying DMN dynamics.

Knowledge of underlying DMN neurophysiology is critical for understanding its dynamic functional properties and relationship to behavior, and for designing network-based treatment regimens for neuropsychiatric and neurological disorders (Greicius et al., 2003; Menon, 2011; Seeley et al., 2007). In particular, computational analyses of causal dynamics in human fMRI data have suggested that behavioral activation of the anterior insular cortex (AI), a key node of the salience network (SN), is implicated in causal deactivation of the DMN (Menon and Uddin, 2010; Nekovarova et al., 2014; Seeley, 2019; Sridharan et al., 2008). However, whether this finding translates to the scale of neuronal activity remains unknown (Menon et al., 2022).

Due to the inherent limitations of noninvasive human fMRI, rodent models are an ideal tool for probing the neural basis and causal underpinnings of DMN dynamics, however the translational utility of these models is currently limited by an incomplete understanding of rodent DMN physiology and function. While there is general agreement that the RSC, and associated posterior medial cortex, anchors the DMN in both rodents and humans (Lu et al., 2012; Sierakowiak et al., 2015), there is uncertainty regarding the differential functional involvement of the Cg and prelimbic cortex (PrL) regions of the rodent mPFC (van Heukelum et al., 2020). In the human brain, the mPFC and the rostral-anterior subdivisions of the Cg are constituents of the DMN (Buckner et al., 2008; Greicius et al., 2003; Raichle, 2015), whereas the dorsal anterior Cg together with the anterior insular cortex (AI) constitute the salience network (SN) (Seeley et al., 2007). Notably, in humans, the SN, and in particular its AI node, plays a critical role in stimulus-driven suppression of the DMN (Menon and Uddin, 2010; Sridharan et al., 2008). This functional segregation of the Cg and mPFC has not been detected in the rodent brain.

Rodent fMRI studies often identify robust co-activations of the RSC, the entirety of Cg, rather than specific sub-regions, and the PrL region of the mPFC; consequently, these areas are all commonly classified as rodent DMN nodes (Belloy et al., 2021; Coletta et al., 2020; Ferrier et al., 2020; Gozzi and Schwarz, 2016; Grandjean et al., 2017, 2020; Gutierrez-Barragan et al., 2019; Hsu et al., 2016; Ji et al., 2018; Lee et al., 2021; Liang et al., 2018; Liska et al., 2015; Lozano-Montes et al., 2020; Lu et al., 2012; Mandino et al., 2022; Nair et al., 2018; Oyarzabal et al., 2022; Paasonen et al., 2018; Peeters et al., 2020a; Stafford et al., 2014; Tu et al., 2021a; Upadhyay et al., 2011; Whitesell et al., 2021; Xu et al., 2022). However, other studies have suggested that PrL and Cg may also be involved in the rodent SN (Gutierrez-Barragan et al., 2019; Mandino et al., 2022; Pagani et al., 2021; Sun and Rebec, 2005; Tsai et al., 2020; Zavala et al., 2003). It follows that to accurately ascribe functional involvement of the Cg and PrL to the rodent SN and DMN it is necessary to further examine the putative nodes of these networks, including the AI, RSC, Cg, and PrL, and investigate their dynamic co-activation and functional connectivity changes.

Critically, the functional organization of the DMN in humans has been characterized not only by synchronization during resting-state conditions, but also by robust deactivation during cognitively demanding tasks (Anticevic et al., 2012; Greicius and Menon, 2004; Greicius et al., 2003; Menon, 2011; Seeley et al., 2007; Shulman et al., 1997). In contrast, the putative rodent DMN has only been characterized under resting-state conditions, and has yet to be validated by this latter, functional definition, thereby presenting a fundamental barrier for translational brain network research. It is therefore crucial to experimentally examine rodent DMN function under awake conditions during attentionally demanding tasks, as in humans.

Here, to form a comprehensive understanding of neuronal signaling among putative rodent DMN and SN areas in a behaviorally relevant context and build an accurate and translational model of the DMN and its dynamic properties, we concurrently measured neuronal calcium activity in the RSC, Cg, PrL, and AI of Thy1-GCaMP6f transgenic rats (Scott et al., 2018). To enable these measurements, we developed an fMRI-compatible, four-channel, spectrally resolved, fiber-photometry recording system based on the platform used in our previous studies (Bao et al., 2020; Chao et al., 2022; Das et al., 2021; Meng et al., 2018; Oyarzabal et al., 2022; Zhang et al., 2022b, 2022a). We used our novel fiber-photometry platform and recent advances in computational modeling of brain-circuit dynamics to characterize neuronal dynamics of the putative DMN nodes and AI under resting-state and awake, salient stimulus presentation conditions.

We first examined neuronal coupling between RSC, Cg, PrL, and AI during the resting-state condition by using time-averaged, static, functional connectivity analysis. We hypothesized that the RSC, Cg, and PrL nodes, belonging to the putative rodent DMN, would form a distinct network. Next, we examined neuronal co-activation patterns associated with simultaneous, fMRI-derived DMN activation and deactivation peaks. We expected to find neuronal antagonism between the AI and the RSC, Cg, and PrL in relation to fMRI-derived DMN activation and deactivation. We then used Multivariate Dynamic State-space systems Identification (MDSI) (Ryali et al., 2011, 2016a, 2016b) to investigate dynamic causal interactions between neuronal activity at the four nodes. Based on previous human fMRI and intracranial electroencephalography (iEEG) studies of causal influences between the AI and DMN (Das and Menon, 2020), we hypothesized that the AI would have a causal role in suppression of the putative rodent DMN nodes. Recent investigations have provided compelling evidence for complex spatiotemporal dynamics associated with the DMN and AI during the resting-state condition in human functional neuroimaging data (Ryali et al., 2016c; Supekar and Menon, 2015). However, as these analyses were based on fMRI, the neuronal basis of these dynamics are unknown. To address this knowledge gap, we employed Bayesian Switching Dynamic Systems (BSDS) state-space algorithms (Taghia et al., 2018) to investigate dynamic changes in neuronal activation and connectivity patterns between the RSC, Cg, PrL, and AI. We hypothesized that the putative DMN nodes (i.e., RSC, Cg, and PrL) would exhibit dynamic coupling and decoupling with the AI.

Finally, we examined the functional organization of the DMN under an awake, freely-moving, condition in response to salient, auditory oddball stimuli – a paradigm that has been shown to consistently activate the AI and deactivate the DMN in human fMRI studies (Crottaz-Herbette and Menon, 2006; Czisch et al., 2012; Labounek et al., 2021; Sridharan et al., 2008). We hypothesized that oddball stimuli would activate the AI and suppress RSC, Cg, and PrL neuronal activity, thereby providing behavioral validation of the putative rodent SN and DMN nodes. By bridging fMRI and neuronal ground-truth recordings with cutting-edge computational analyses, we were able to uncover several novel dynamic properties of DMN and SN interactions in the rodent brain and establish the neuronal underpinnings of their antagonistic relationship commonly observed with hemodynamic fMRI techniques. Our findings advance foundational knowledge of the neuro-functional organization of the rodent DMN, and provide a more informed model for translational studies.

## Results

### Time-averaged, static, functional connectivity in AI, Cg, PrL, and RSC neuronal GCaMP, resting-state signals

We used an fMRI-compatible, spectrally resolved, fiber-photometry platform (Bao et al., 2020; Chao et al., 2022; Das et al., 2021; Meng et al., 2018; Oyarzabal et al., 2022; Zhang et al., 2022b, 2022a) with four recording fibers (**Figure 1A**) to measure neuronal GCAMP activity in the RSC, Cg, PrL, and AI from rats (**Supplementary Figure 1**). GCaMP activity was combined with whole-brain fMRI recordings in anesthetized rodents to capture concurrent resting-state signals from both modalities (**Figure 1A**). First, we investigated the functional coupling of neuronal signals among the four targeted brain regions. Time-frequency analysis of neuronal GCaMP data in the RSC, Cg, PrL, and AI revealed prominent low-frequency spectral power fluctuations (**Figure 1B-E**). We calculated time-averaged, static, functional connectivity of the GCaMP signals and revealed significant functional connectivity between the RSC and Cg, between the Cg and PrL, and between the AI and PrL (**Figure 1F**, *p* < 0.01, two-tailed *t*-test, FDR-corrected). We then used hierarchical clustering to examine the similarity of functional connectivity profiles. This analysis revealed that the RSC has the strongest link with the Cg, a weaker link with the PrL, and the weakest link with AI (**Figure 1G**). These results demonstrate hierarchical organization of functional connectivity between putative rodent DMN nodes and AI at the neuronal level.

**Figure 1.**
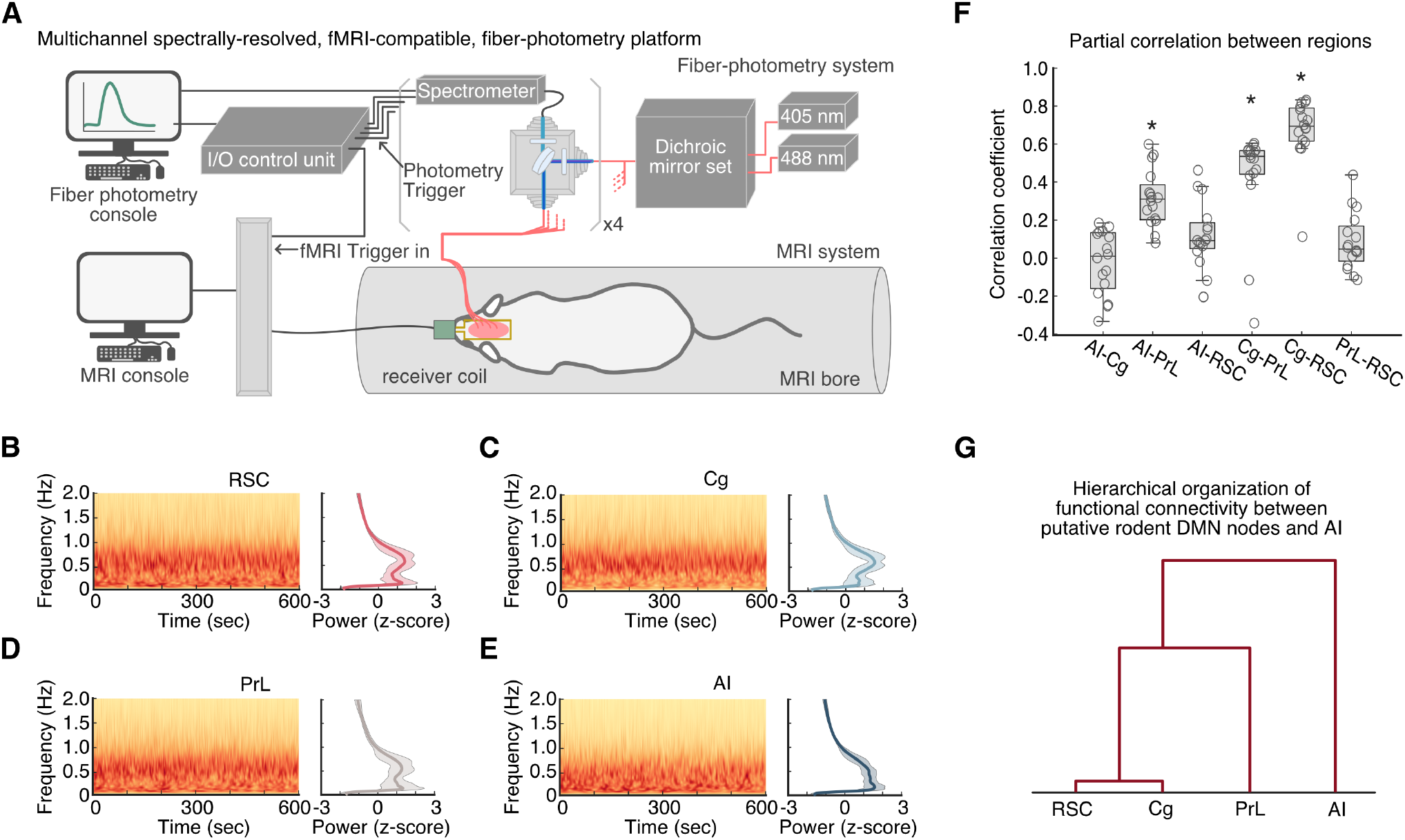
A multichannel, fMRI-compatible, spectrally-resolved, fiber-photometry platform used to measure coupling of neuronal signals between the AI, Cg, PrL, and RSC. (**A**) Schematic illustration of the photometry system used for measuring GCaMP signals in four distinct brain areas concurrently with fMRI. **(B)** Brain regions targeted by optical fibers for photometry recording of neuronal GCaMP activity. **(B-E)** Spontaneous, resting-state, GCaMP signal fluctuations in AI, Cg, PrL, and RSC in anesthetized rats. Representative GCaMP time-frequency plots and corresponding average GCaMP power spectra were plotted for each region, respectively. Prominent spectral power below 1Hz was found in all photometry recording sites. **(F)** Partial correlations between GCaMP activity recorded from the AI, Cg, PrL, and RSC included significant positive correlations between the AI and Cg, the Cg and PrL, and the Cg and RSC. Data are presented as box and whisker plots, where boxes encompass values between the 25^th^ and 75^th^ percentiles and horizontal lines represent median values. Dots in the figure represent the correlation coefficients of each individual rat. *\*p* < 0.01, two-tailed *t*-test, FDR-corrected. **(G)** Dendrogram from hierarchical analysis of the strength of functional connectivity shown in **F**.

### Resting-state neuronal GCaMP dynamics in relation to fMRI-derived DMN activation and deactivation

We next characterized neuronal signals associated with concurrent fMRI-derived DMN activation and deactivation peaks. We used brain-wide CBV-fMRI signals (Kim et al., 2013) with group-level independent component analysis (gICA) (Smith et al., 2004) and dual regression (Zuo et al., 2010) to identify the DMN **(Figure 2A**) and its associated time series **(Figure 2B, top**), similar to previous rat DMN fMRI studies (Hsu et al., 2016; Liu et al., 2016; Ryali et al., 2016a; Wen et al., 2013). Individual-level DMN time-courses from the dual regression procedure were used to identify DMN activation and deactivation events corresponding to local maxima or minima (*z* > 1.64, *p* < 0.05) **(Figure 2B, top)**. We then extracted neuronal GCaMP signals from the RSC, Cg, PrL and AI and examined tlme-locked responses in these signals associated with activation and deactivation peaks in the fMRI-derived DMN time series (**Figure 2B, bottom**). Our analysis revealed robust changes in neuronal GCaMP signals time-locked to the fMRI-derived DMN activation and deactivation peaks (**Figure 2C and D**). Notably, the RSC, Cg and PrL showed complex neuronal GCaMP signal changes (increased power at roughly 0.5 Hz followed by decreased power at around 1 Hz) while the AI showed decreased power at low frequency band preceding the fMRI-derived DMN activation peaks (**Figure 2C**). Neuronal GCaMP signal changes in all four regions preceded DMN activation by 2-6 seconds (**Figure 2C**, **Supplementary Figure 2**). Further, neuronal GCaMP signal power at the low frequency band decreased in the RSC but increased in the AI preceding the fMRI-derived DMN deactivation peaks (**Figure 2D**). These results demonstrate similar time-locked patterns in the RSC, Cg and PrL, and reveal neuronal antagonism between the AI and the three other brain regions in relation to fMRI-derived DMN activation and deactivation.

**Figure 2.**
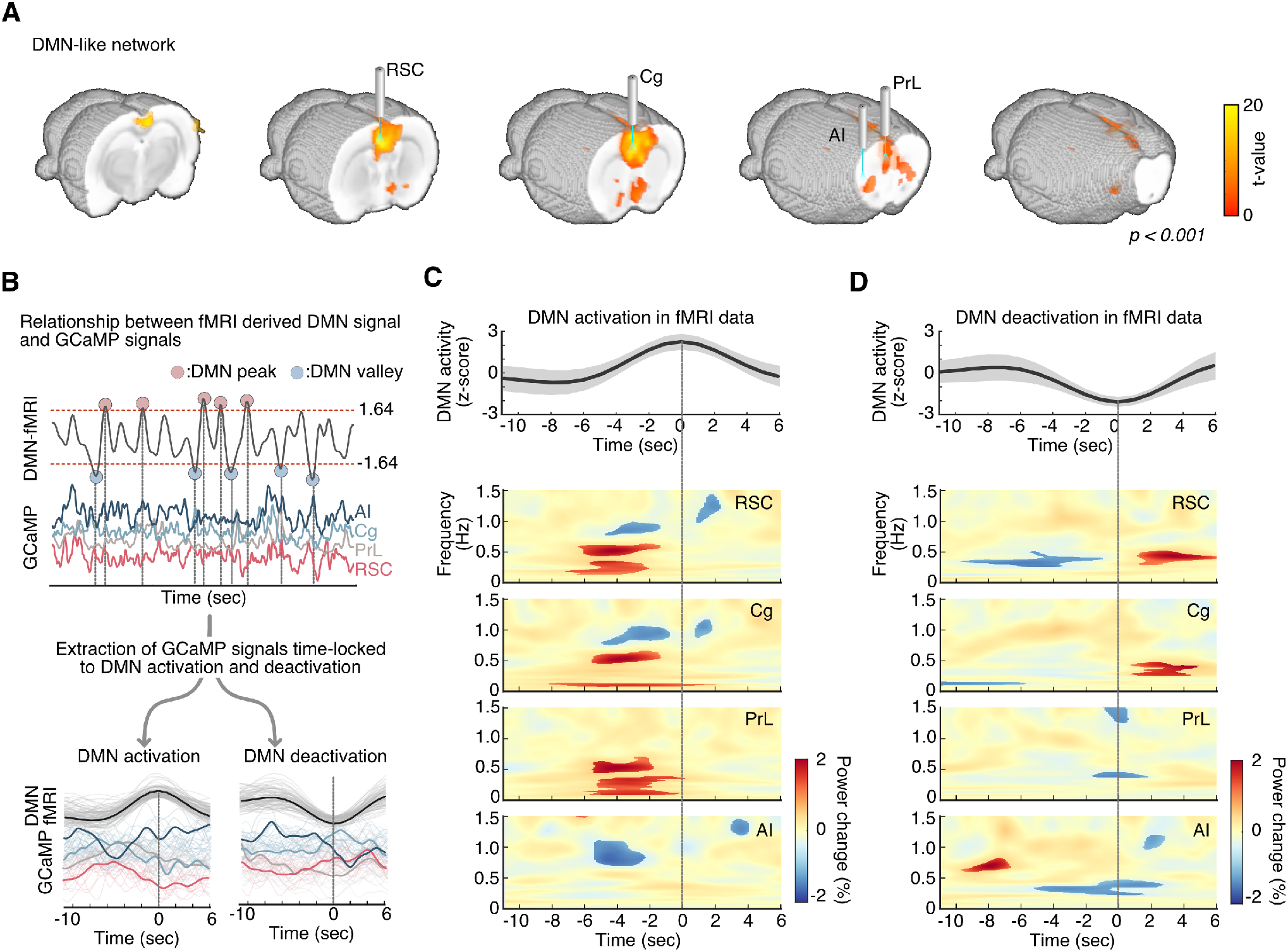
Frequency-specific GCaMP power changes associated with concurrent fMRI-derived DMN activations and deactivations. **(A)** Brain regions corresponding to DMN-related nodes targeted by optical fibers for photometry recording of GCaMP and CBV-fMRI-derived DMN map (one sample *t*-test, *t*-value > 4, *p* < 0.001). **(B)** Schematic illustrating identification of AI, Cg, PrL and RSC neuronal activity time-locked to DMN activation and deactivation. Subject-level DMN time series were used to identify DMN activation and deactivation events (local maximums z < 1.64, local minimums z < 1.64, *p* < 0.05). **(C)** Top: Averaged DMN activation peak in fMRI data. Bottom: Statistical maps of GCaMP spectrograms from each node time-locked to DMN activation peaks (*p* < 0.05, two-tailed *t*-test). **(D)** Top: Averaged DMN deactivation peak in fMRI data. Bottom: Statistical maps of GCaMP spectrograms from each node time-locked to DMN deactivation peaks (*p* < 0.05, two-tailed *t*-test).

### Causal interactions between RSC, Cg, PrL, and AI resting-state, neuronal GCaMP signals

Next, we used neuronal GCaMP recordings to test the hypothesis that the AI has a causal role in the suppression of DMN nodes. We applied MDSI algorithms (Ryali et al., 2011) on time series data from the four regions. This analysis revealed inhibitory, causal outflow from the AI to the RSC, Cg, and PrL (**Figure 3**, *p* < 0.01, two-tailed *t*-test, FDR-corrected). These results identify a circuit mechanism underlying neuronal antagonism between the AI and the RSC, Cg, and PrL.

**Figure 3.**
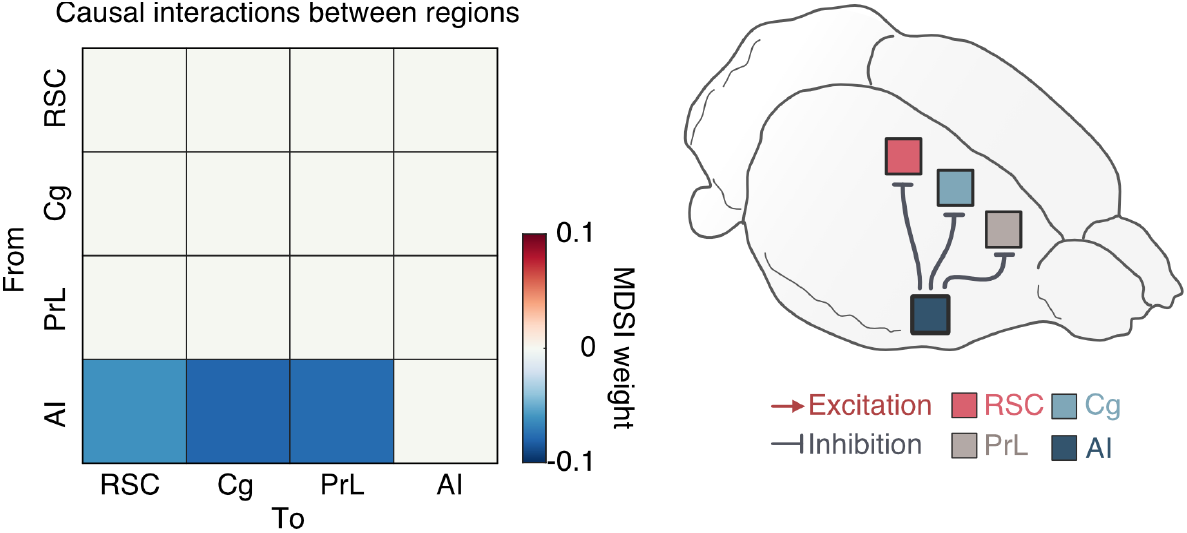
Causal interactions between the AI, Cg, PrL, and RSC during the resting-state condition in anesthetized rats. Causal interactions between regions were identified by using multivariate dynamical systems identification (MDSI) (Ryali et al., 2011). In the resting-state, MDSI showed significant inhibitory causal influence from the AI to the Cg, PrL, and RSC in the rat brain (*p* < 0.01, FDR-corrected).

### Dynamic brain states associated with RSC, Cg, PrL and AI resting-state, neuronal GCaMP signal interactions

We next investigated time-varying spatiotemporal dynamics associated with the DMN and AI during the resting-state condition. We used BSDS, a probabilistic state-space model for automatically identifying latent brain states, their time-evolving patterns of activation and connectivity patterns, occupancy rates, and state transition probabilities (Taghia et al., 2018). In each rat, BSDS estimated the posterior probability of each latent brain state at each time point (**Figure 4A, top**), and the brain state with the highest posterior probability was chosen as the dominant state at that time point for that rat (**Figure 4A, bottom**).

**Figure 4.**
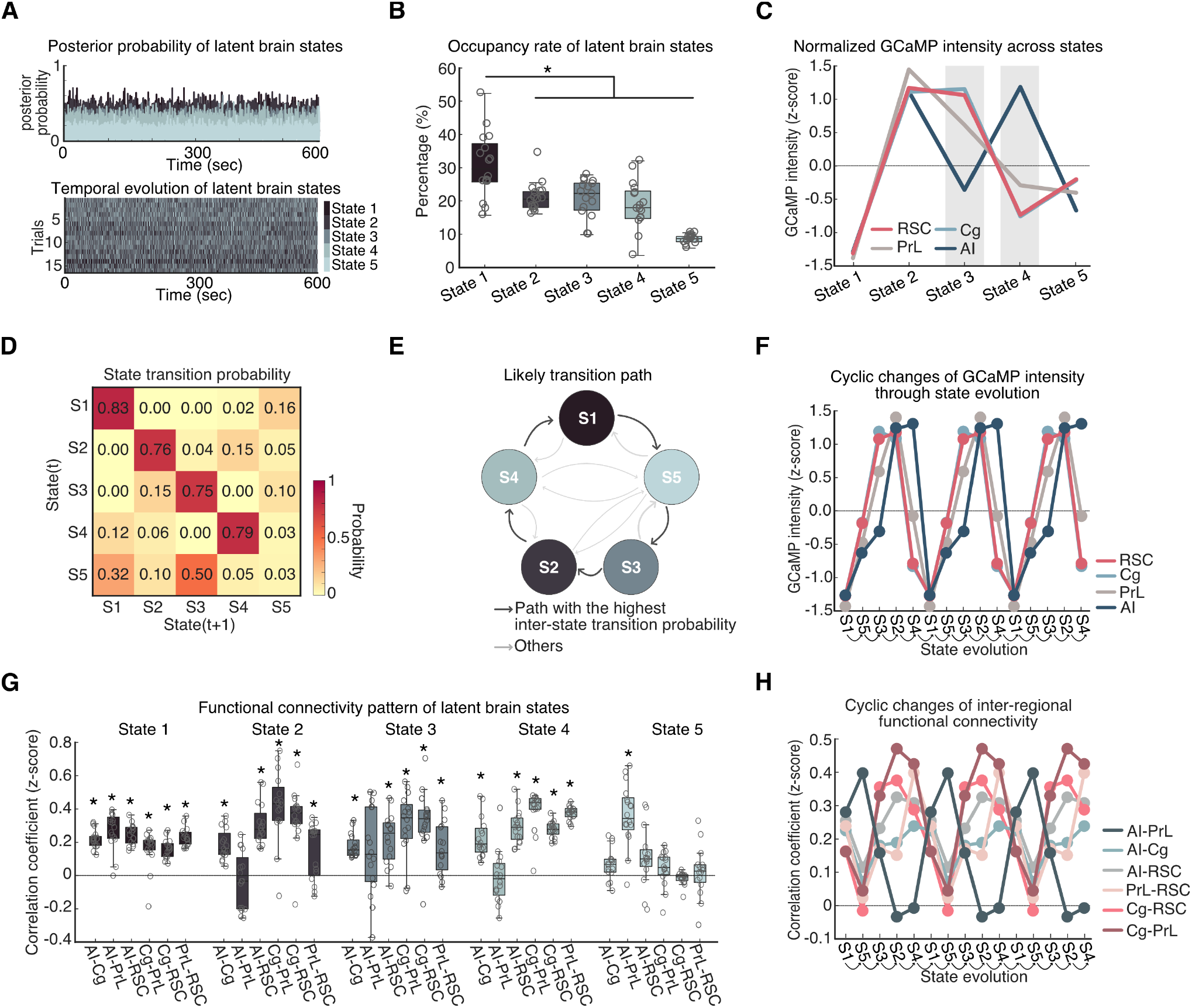
Spatiotemporal properties of latent brain states during the resting-state condition. **(A)** Top: Time-varying posterior probabilities of the brain states identified by the Bayesian switching dynamical systems (BSDS) model. Bottom: Temporal evolution of the identified brain states. **(B)** Occupancy rates of the latent brain states. The occupancy rate of State 1 was significantly higher than other brain states. Data are presented as box and whisker plots, where boxes encompass values between the 25^th^ and 75^th^ percentiles and horizontal lines represent median values. Dots in the figure represent correlation coefficients for each individual rats. \**p* < 0.01, two-tailed *t*-test, FDR-corrected. **(C)** Normalized GCaMP activities of AI, Cg, PrL, and RSC across latent brain states. Cg, PrL, and RSC showed coupled activities across all the states whereas the activity of AI showed coupling and decoupling with activities of Cg, PrL and RSC. **(D)** Dynamic switching properties of latent brain states. State switching probability, defined as the probability that a latent brain state at time instance (*t*) stays in its own state or switches to other states at time instance (*t* + *1*). **(E)** Analysis of likely switching paths revealed that state transition follows a cyclic pattern. **(F)** Evolution of GCaMP activities of four regions following the likely transition path identified in **Figure 4E**. Evolution of GCaMP activity exhibits a cyclic pattern with coupling and decoupling between AI and other regions. **(G)** Functional connectivity pattern between regions for each latent brain state. Data are presented as box and whisker plots, where boxes encompass values between the 25^th^ and 75^th^ percentiles and horizontal lines represent median values. Dots in the figure represent the correlation coefficients from each individual rat. \**p* < 0.01, two-tailed *t*-test, FDR-corrected. **(H)** Evolution of inter-regional functional connectivity (Fisher Z-transformed partial correlation coefficient) following the likely transition path identified in **Figure 4E**.

BSDS identified five brain states, each characterized by unique spatiotemporal dynamics. We determined the occupancy rate of each brain state, which quantifies the fraction of time that a given state is most likely to occur. This analysis revealed that State 1 had a significantly higher occupancy rate (30.6±2.5%, Mean±SEM) than other states (all *p* < 0.01, two-tailed *t*-test, FDR-corrected, **Figure 4B**). The occupancy rates of States 2, 3, and 4 were each around 20%, respectively (State 2: 21.1 ± 1.1%, State 3: 20.9±1.4%, State 4: 18.8±1.8%), while State 5 had the lowest (8.5±1.3%). These results demonstrate that, at the neuronal level, interactions between the RSC, Cg, PrL and AI are characterized by multiple dynamically evolving brain states rather than a single, static state, consistent with observations from fMRI studies (Ryali et al., 2016c).

### Distinct state-specific activation and deactivation patterns in resting-state, neuronal GCaMP signals

We next examined activation levels in the RSC, Cg, PrL and AI associated with each brain state estimated by BSDS. This analysis revealed distinct activation patterns across the four regions in each brain state (**Figure 4C**). All four regions were deactivated during State 1, with neuronal GCaMP signals well below the average across all states. A similar, but subtler deactivation across all regions was also observed in State 5. In contrast, neuronal GCaMP signals were all above average during State 2. In States 3 and 4, the RSC, Cg, and PrL showed different activation patterns than the AI. Specifically, in State 3, activation was above average in the RSC, CG, and PrL, but below average in the AI. In State 4, this pattern was reversed with RSC, CG, and PrL below average and the AI above average. These results demonstrate that resting-state neuronal activity is characterized by similar patterns of co-activation and co-deactivation among the RSC, Cg, PrL and AI in States 1, 2, and 5 which occur about 60% of the time. However, for the remaining 40% of the time, the AI showed a dissociable pattern of activation and deactivation from the three other brain areas. These five brain states and their activation/deactivation patterns were also identifiable by k-means clustering (**Supplementary Figure 3**), supporting the reliability of the observed brain state patterns.

### Dynamic state transitions during the resting-state condition

Next, we investigated dynamic transition properties between the distinct brain states identified in neuronal GCaMP resting-state data. We used BSDS-derived state transition probabilities for each rat and determined the most likely transition path between brain states. The state transition matrix also indexes the stability of brain states by calculating the probability that a brain state remains the same from a time point *t* to the next time point *t+1*. Analysis of state transition probabilities revealed that States 1, 2, 3, and 4 were not volatile from one-step to another, but instead persisted over time (*p* < 0.001, two-tailed *t*-test, FDR-corrected; **Figure 4D**). State 5, however, was highly volatile (*p*>0.05, two-tailed *t*-test). Our analysis further revealed a canonical transition path from S1 to the other states and back again: S1➔S5➔S3➔S2➔S4➔S1 **(Figure 4D and E**, **Supplementary Figure 4**). Notably, examination of the evolution of brain states associated with dynamic GCaMP neuronal activity revealed a cyclical pattern of activation and deactivation involving the RSC, Cg, PrL and AI (**Figure 4F**). These results demonstrate that state transitions do not occur in random order, but rather follow specific transition patterns.

### State-dependent changes in resting-state functional connectivity

We then used results from the BSDS state-space model to investigate changes in functional connectivity across states. Examination of functional connectivity revealed distinct connectivity patterns associated with each state (**Figure 4G**). State 1 was characterized by significant positive functional connectivity among all regions (*p* < 0.01, two-tailed *t*-test, FDR-corrected). States 2, 3, 4 showed significant positive functional connectivity between all of the regions except between the AI and PrL, whereas State 5 showed significant positive functional connectivity between the AI and PrL but not between other links (*p* < 0.01, two-tailed *t*-test, FDR-corrected).

Examination of functional connectivity associated with the most likely transition path (identified in **Figure 4E)** revealed a cyclical pattern of changes characterized by synchronized activity among all pairs of regions except between the AI and PrL (**Figure 4H**). Specifically, during the S1➔S5➔S3➔S2➔S4➔S1 cycle, the PrL showed the highest synchronization with AI and lower synchronization with Cg and RSC during S1 and S5, but then exhibited the lowest synchronization with AI and higher synchronization with Cg and RSC during S3, S2, and S4. The engagement of PrL with not only other putative DMN nodes, but the AI as well, suggests that PrL may have a dual-role in both the DMN and the SN.

### Auditory oddball stimuli-induced AI, Cg, PrL, and RSC neuronal GCaMP responses in awake, freely moving rats

Thus far, rodent studies have mostly assigned nodes to the DMN based on synchronous resting-state activity without assessing the functional responses of these nodes to salient external events. Therefore, it is crucial to validate putative rodent DMN nodes in a behaviorally relevant context (Menon and Uddin, 2010; Sridharan et al., 2008). To this end, we recorded neuronal GCaMP signals from the AI, PrL, Cg, and RSC (*vide supra*) of awake, freely moving rats during an auditory oddball paradigm – a paradigm that has been shown to consistently activate the AI and deactivate the DMN in human fMRI studies (Czisch et al., 2012; Labounek et al., 2021; Sridharan et al., 2008). Specifically, rats were presented with standard and deviant (i.e., oddball) tones with occurrence rates of 97% and 3%, respectively (**Figure 5A**). Peri-event time-frequency analysis of neuronal GCaMP signals revealed significant deactivation of the RSC, Cg, and PrL (0.5-8 seconds post-oddball stimuli) and activation of the AI (2-3 seconds post-oddball stimuli) following oddball stimuli (*p* < 0.05, two-tailed *t*-test; **Figure 5B**). These results demonstrate that the RSC, Cg, and PrL are deactivated by salient stimuli, thus providing functional evidence for their classification as DMN nodes.

**Figure 5.**
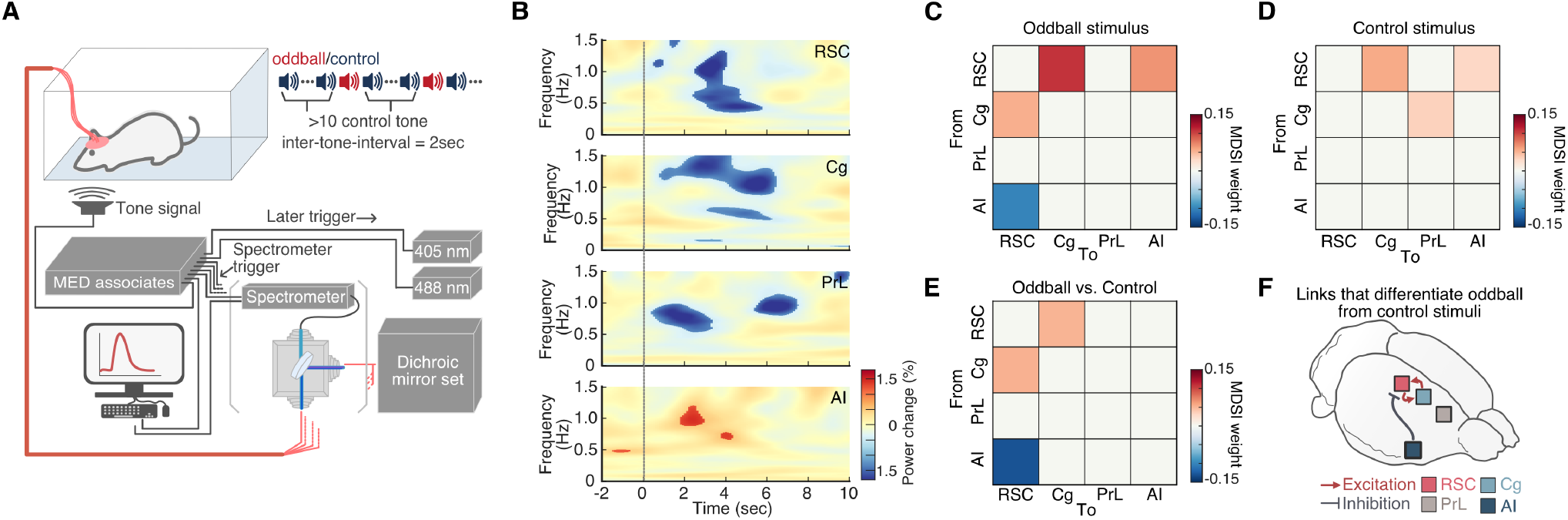
Neuronal activity changes time-locked to acoustic oddball stimuli. **(A)** Schematic illustrating the oddball experiment paradigm. **(B)** Statistical map of GCaMP spectrogram (*p* < 0.05, two-tailed *t*-test) and percentage GCaMP power changes time-locked to acoustic oddball stimuli presentations (vertical line). **(C-F)** Causal interactions between the AI, Cg, PrL, and RSC during oddball stimulation. Causal relationships between regions were estimated during oddball and control stimuli presentations using MDSI. **(C)** During oddball stimuli, significant inhibitory causal influence was observed from the AI to the RSC, and significant excitatory causal influence was observed from the RSC to the Cg and AI and from the Cg to the RSC (all *p* < 0.01, FDR-corrected). **(D)** Significant excitatory causal influence was observed from the RSC to the Cg and AI and from the Cg to the PrL during control stimuli (all *p* < 0.01, FDR-corrected). **(E)** Compared to control stimuli, oddball stimuli evoked significantly stronger inhibitory causal influence from the AI to the RSC and significantly stronger bidirectional excitatory influences between the Cg and the RSC (all *p* < 0.01, two-tailed *t*-test, FDR-corrected). **(F)** Summary illustration of the specific links that showed significant differences in causal interactions between oddball and control stimuli.

### Causal interactions between the AI, Cg, PrL, and RSC neuronal GCaMP signals during the auditory oddball paradigm in awake, freely moving rats

We next examined causal interactions between the AI, Cg, PrL, and RSC neuronal GCaMP signals during the oddball paradigm using MDSI. Based on the activation profile of the AI in **Figure 5B**, we used GCaMP signals from 4 seconds post-oddball stimuli to investigate causal interactions related to oddball stimuli. This analysis identified significant inhibitory causal influence from the AI to the RSC and excitatory causal influence from the RSC to the Cg and AI, and from the Cg to the PrL during processing of oddball stimuli (*p* < 0.01, two-tailed *t*-test, FDR-corrected; **Figure 5C**). In comparison to frequent control stimuli, oddball stimuli produced stronger inhibitory causal influence from the AI to the RSC, and additional bidirectional excitatory influences between the Cg and the RSC (*p* < 0.01, two-tailed *t*-test, FDR-corrected; **Figure 5D-F**). These results provide neuronal evidence from GCaMP recordings that the AI suppresses the DMN via inhibitory control over the RSC in awake, behaving rodents.

## Discussion

We investigated the neuronal origins of the rodent DMN using GCaMP recordings from resting-state and auditory oddball task conditions. We used transgenic rats expressing genetically encoded calcium indicators under the Thy1 promoter, allowing direct measurement of pyramidal neuron activity using an fMRI-compatible, four-channel, spectrally resolved fiber-photometry platform. We examined dynamic activation and connectivity from GCaMP recordings in DMN- and SN-related regions in the rodent brain. Our findings reveal: **(1)** robust neuronal resting-state connectivity between the RSC, Cg, and PrL; **(2)** neuronal antagonism between the AI and the RSC, Cg, and PrL in relation to fMRI-derived DMN activation and deactivation peaks; **(3)** significant neuronal inhibitory causal outflow from the AI to the RSC, Cg, and PrL during the resting-state condition; **(4)** cyclical state transitions characterized by the neuronal activity in AI being intermittently in- and out-of-phase with activity of RSC, Cg, and PrL during the resting-state condition; **(5)** cyclical changes in inter-regional connectivity characterized by a unique pattern of the dynamic synchronization between PrL and AI that was out-of-phase from all other pairs of regions; **(6)** salient stimulus-induced activation of AI and deactivation of the RSC, Cg and PrL in awake, freely moving rats; and **(7)** salient stimulus-induced dynamic causal suppression of the RSC by the AI in awake, freely moving rats. Together, these results provide neuronal GCaMP-based evidence that the RSC, Cg and PrL function as DMN nodes in the rodent brain, with the RSC and Cg showing the closest correspondence in their activity and connectivity profiles and the PrL exhibiting a potentially dual-purpose role at the interface between the DMN and SN. Our results also reveal a neuronal basis of causal inhibitory control signals from the AI to the RSC that facilitate access to attentional resources as observed in hemodynamic fMRI studies (Menon and Uddin, 2010; Sridharan et al., 2008). Importantly, the existence of causal inhibitory control of the DMN by the AI in the rodent brain highlights the utility of rodent models in probing the cellular and circuit mechanisms that control network switching during behavior.

In humans, the DMN has been linked to self-referential functions involving autobiographical memory, recollection, and planning (Binder et al., 1999; Buckner et al., 2008; Greicius et al., 2003; Mars et al., 2012; Spreng et al., 2009), and abnormal DMN activity and connectivity have been reported in many neurological and neuropsychiatric disorders, including Alzheimer’s disease (Greicius et al., 2004), Parkinson’s disease (Hou et al., 2017; Tessitore et al., 2012), temporal lobe epilepsy (Liao et al., 2011; Zhang et al., 2010), attention deficit hyperactivity disorder (Uddin et al., 2008, 2010; Wilson et al., 2013), and mood disorders (Liston et al., 2014; Sheline et al., 2009). Due to its substantial importance in basic and clinical science, significant translational efforts have been made to examine the functional organization of a putative DMN in rodent brains. In this study, using a novel MRI-compatible, multichannel, spectrally resolved fiber-photometry platform, we identify the neurofunctional organization of the rodent DMN and provide a more informed model for translational studies using the plethora of biological and genetic tools now available for neural circuit dissection in rodent models.

The first goal of our study was to investigate intrinsic neuronal coupling between putative rodent DMN nodes in the resting state condition. While studies using resting-state fMRI have suggested that the rat DMN is comprised of the RSC, Cg, and PrL (Gozzi and Schwarz, 2016; Hsu et al., 2016; Ji et al., 2018; Lee et al., 2021; Liang et al., 2018; Lu et al., 2012; Osmanski et al., 2014; Oyarzabal et al., 2022; Paasonen et al., 2018; Peeters et al., 2020a; Tu et al., 2021b, 2021a; Xu et al., 2022), such coupling has not yet been demonstrated at the neuronal scale. Our findings using static time-averaged functional connectivity analysis are in general agreement with prior literature, showing strong coupling between RSC, Cg, and PrL in the anesthetized rat brain (Gozzi and Schwarz, 2016; Hsu et al., 2016; Ji et al., 2018; Lee et al., 2021; Liang et al., 2018; Lu et al., 2012; Osmanski et al., 2014; Oyarzabal et al., 2022; Paasonen et al., 2018; Peeters et al., 2020a; Tu et al., 2021b, 2021a; Xu et al., 2022), as well as anesthetized mouse brains (Belloy et al., 2021; Coletta et al., 2020; Ferrier et al., 2020; Gozzi and Schwarz, 2016; Grandjean et al., 2017, 2020; Gutierrez-Barragan et al., 2019; Liska et al., 2015; Mandino et al., 2022; Stafford et al., 2014; Vanni et al., 2017; Whitesell et al., 2021). Additionally, our analysis revealed that RSC, Cg and PrL exhibit similar time-locked neuronal activity patterns while the AI displays neuronal antagonism with these regions in relation to fMRI-derived DMN activation and deactivation. Further, causal circuit analysis revealed inhibitory causal outflow from the AI to the RSC, Cg, and PrL, providing a more precise circuit mechanism underlying neuronal antagonism between AI and RSC, Cg, and PrL. Taken together, these findings identify key aspects of the intrinsic functional organization of the rodent DMN at the neuronal level and suggest that the widely documented AI-DMN antagonistic relationship in the human brain (Menon, 2011; Menon and Uddin, 2010; Nekovarova et al., 2014; Seeley, 2019; Sridharan et al., 2008) also exists in the rat brain. Given the wide range of cellular manipulation and recording techniques available in rodent species, our findings open a vast avenue for further investigations of large-scale brain networks, including the DMN and SN.

The next goal of our study was to identify dynamic changes in interactions between putative rodent DMN nodes using neuronal recordings and state-space modeling, building on our prior work in humans (Ryali et al., 2016c). While there is broad agreement that the RSC anchors the rodent DMN similarly to the human DMN, there is less consensus about the relative roles of the Cg and PrL. Although many studies have suggested that the Cg and PrL are part of the rodent DMN (Belloy et al., 2021; Coletta et al., 2020; Ferrier et al., 2020; Gozzi and Schwarz, 2016; Grandjean et al., 2017, 2020; Gutierrez-Barragan et al., 2019; Hsu et al., 2016; Ji et al., 2018; Lee et al., 2021; Liang et al., 2018; Liska et al., 2015; Lozano-Montes et al., 2020; Lu et al., 2012; Mandino et al., 2022; Nair et al., 2018; Oyarzabal et al., 2022; Paasonen et al., 2018; Peeters et al., 2020a; Stafford et al., 2014; Tu et al., 2021a; Upadhyay et al., 2011; Whitesell et al., 2021; Xu et al., 2022), other resting-state fMRI studies have shown that the Cg and PrL may also form part of the rodent SN (Gutierrez-Barragan et al., 2019; Mandino et al., 2022; Pagani et al., 2021; Sun and Rebec, 2005; Tsai et al., 2020; Zavala et al., 2003). Crucially, hierarchical clustering of static functional connectivity patterns revealed that the RSC has the strongest link with the Cg, a weaker link with the PrL, and the weakest link with AI.

It is plausible that the rodent mPFC, which is less-spatially-segregated than in humans, is functionally engaged in both the DMN and SN. To shed light on the potentially multiplexed and dynamic involvement of the Cg and PrL of the mPFC in the rodent DMN and SN and enable modeling of dynamic state changes in activity and connectivity patterns among the RSC, Cg, PrL, and AI, we employed multi-region neuronal GCaMP activity recordings, with superior sampling rate and signal kinetics than conventional fMRI. State-space modelling revealed synchronization of the RSC, Cg and PrL GCaMP signals across all states (states 1-5), which suggests a resilient neuronal functional network formed by these nodes in the resting rodent brain and provides a neuronal foundation for the fMRI-derived DMN. Consistent with our findings, pharmacological activation of the PrL (Saitoh et al., 2014; Suzuki et al., 2016), or enhancement of Cg activity through basal forebrain modulation (Lozano-Montes et al., 2020; Nair et al., 2018), has been shown to promote self-directed behaviors associated with the DMN. Chemogenetic inhibition of Cg also reduces Cg-RSC functional connectivity (Peeters et al., 2020b) and suppresses low frequency power of fMRI signals in the PrL and RSC (Tu et al., 2021a). Likewise, chemogenetically enhanced low frequency power of neuronal activity in the PrL strengthens DMN connectivity in mice (Rocchi et al., 2022). Increased neuronal spiking in the Cg has also been associated with strengthened functional connectivity within the frontal DMN nodes, as well as between frontal and association cortex DMN nodes in mice (Oyarzabal et al., 2022).

Our analysis also revealed that AI exhibits both dynamic coupling (State 1, 2, 5) and decoupling (state 3 and 4) from the RSC, Cg and PrL suggesting that the AI dissociates during the emergence of DMN co-activation patterns. Such dynamic decoupling of the AI from these DMN nodes is similar to what has been observed in human fMRI (Menon, 2011; Menon and Uddin, 2010; Nekovarova et al., 2014; Seeley, 2019; Sridharan et al., 2008), and lends further support for the roles of both the Cg and PrL in the DMN. Notably, examination of functional connectivity associated with the most-likely transition path (**Figure 4E)** revealed a consistent cyclical pattern of changes, in which the AI-PrL synchronization was out of phase from all other pairs of regions (**Figure 4H**). Specifically, PrL activity was synchronized with the RSC and Cg during States 2, 3 and 4, and switched to higher levels of synchronization with the AI during States 1 and 5. These intriguing findings suggest that in addition to the DMN, the PrL may also play a transient dual-role in the SN. These findings are convergent with and clarify the dynamic underpinnings of hierarchical connectivity patterns observed with the static functional connectivity analysis (**Figure 1G**). Furthermore, our findings identify dynamic switching of AI-PrL synchrony as a prominent feature underlying the transition of brain networks.

The final goal of our study was to characterize the functional modulation of neuronal activity in rodent DMN nodes and the AI by salient stimuli. As noted, the functional organization of the human DMN has been identified not only by synchronous activity at rest, but also by consistent patterns of suppression during attention to salient environmental events (Greicius et al., 2003; Menon and Uddin, 2010; Raichle et al., 2001). Rodent studies, however, have mostly assigned nodes to the DMN based on synchronous resting-state activity without assessing their functional response to salient events in the awake condition. It is therefore crucial to characterize putative rodent DMN nodes in a behaviorally relevant context (Menon and Uddin, 2010; Sridharan et al., 2008). Although multiple nodes of the rodent DMN has not been investigated in awake, behaving conditions, a few studies have examined the RSC node of the DMN, specifically, under those conditions. Ferrier et al. observed whisker-stimulation-induced blood volume reduction in the RSC using single-slice functional ultrasound imaging in the mouse brain (Ferrier et al., 2020), and Fakhraei et al. reported suppression of local field potentials in RSC during an externally oriented visual task in the rat brain (Fakhraei et al., 2021). Our study fills a critical gap by concurrently measuring neuronal activity from the three major putative DMN nodes – RSC, Cg, and PrL. Importantly, we identified concurrent deactivation of the RSC, Cg and PrL in response to salient stimuli, providing strong support for their functional involvement in the rodent DMN. We also identified activation of the AI in response to salient stimuli in awake, freely moving rats, confirming its role in encoding salient stimuli established in human fMRI studies (Menon and Uddin, 2010; Sridharan et al., 2008). Crucially, in an advance over previous fMRI studies, we also found that the AI has a strong causal influence on the RSC and its suppression during the processing of salient oddball stimuli. Our findings have critical implications for the study of the precise causal role of the AI, RSC, Cg, and PrL in behavior and large-scale brain network switching, and demonstrate that casual influences between the AI and RSC are not unique to hemodynamic fMRI signals.

## Summary and conclusions

We elucidate the functional properties of critical DMN and SN nodes in the rat brain using a novel neuronal GCaMP and fMRI recording platform. We found that neuronal activity in the RSC, Cg and PrL are co-activated under the resting-state condition. Our causal circuit analysis demonstrated significant inhibitory neuronal causal outflow from the AI to the RSC, Cg and PrL, which parallel human fMRI findings, and provide neuronal evidence that the AI plays a crucial role in dynamic network switching (Sridharan et al., 2008). Our results include neuronal activity-based evidence of the RSC, Cg and PrL forming a single hierarchical network, representing the rat DMN. Our analysis revealed cyclical state transitions during the resting-state condition characterized by dynamic switching between DMN nodes and the AI, with the AI intermittently in- and out-of-phase with DMN nodes. We also identified changes in AI and PrL activity synchronization over distinct phases of brain state evolution that were out of phase with the connectivity between other node-pairs. We further established a functional homology between rat and human DMN and SN using an auditory oddball task in awake free-moving rats. We found that salient oddball stimuli activate AI while suppressing neuronal activity in the RSC, Cg and PrL. Lastly, causal circuit analysis revealed salient-stimuli-induced, dynamic, causal suppression of the RSC by the AI, and causal co-activation between Cg and RSC. These findings identify a translational and neural correspondence between AI and DMN responses in rats to salient task-relevant stimuli, and show that the rodent AI plays a causal role in DMN-SN dynamic switching. Together, our work paves the way for future translational studies using rodent models to investigate the cellular basis of cognitive control circuits, such knowledge can help understand the dynamic properties of brain states, define their relationship to behaviors, and ultimately design network-based treatment regimens for neuropsychiatric and neurological disorders.

## Acknowledgements

This work is supported in part by NIH grants R01MH126518, RF1NS086085, RF1MH117053, R01MH111429, R01NS091236, P60AA011605, P50HD103573, S10MH124745 and S10OD026796.

## Author contributions

T.-H.H.C., L.-M.H., D.H.C., V.M., and Y.-Y.I.S. conceived the project and designed the experiments. T.-H.H.C., W.-T.Z., and G.C. implemented the methods. T.-H.H.C. executed experiments. T.-W.W.W. performed histology. T.-H.H.C., L.-M.H., and B.W.L. analyzed data. T.-H.H.C., D.H.C., L.-M.H., B.W.L., V.M., and Y.-Y.I.S. wrote the manuscript with input from all authors. V.M. and Y.-Y.I.S. supervised the study and provided funding support.

## Declaration of interests

The authors declare no competing interests.

## Methods

### KEY RESOURCES TABLE

| REAGENT OR RESOURCE | SOURCE | IDENTIFIER |
| --- | --- | --- |
| Software and Algorithms |  |  |
| MATLAB | Mathworks | R2019b |
| Other |  |  |
| 405 nm laser and remote | Coherent | OBIS 405 LS (100 mW) |
| 488 nm laser and remote | Coherent | OBIS 488 LS (100 mW) |
| Spectrometer | Ocean Optics | QE-pro |
| Optic patch cable | Thorlabs | FT200EMT (200 mm, 0.39 NA) |
| Uncleaved fiber optic cannula | Thorlabs | CFMC12U-20 (200 mm, 0.39 NA) |

#### Stereotactic surgery and animal preparation

We used Thy1-GCaMP6f transgenic rats (Scott et al., 2018) which have the fluorescent calcium activity indicator, GCaMP (Chen et al., 2013), expressed under Thy1 promoter (Feng et al., 2000), allowing measurement of cortical output activity from pyramidal neurons. The rats were initially anesthetized by 5% isoflurane, then maintained under anesthesia by a constant flow of 2-2.5% isoflurane mixed with medical air. Rectal temperature was continuously monitored and maintained within 37 ± 0.5 °C. Rats were positioned on a stereotactic frame (Model 962, Kopf Instruments) with ear bars and a tooth bar. The scalp was removed to expose the skull (~1×1 cm). Burr holes were prepared according to experimental coordinates, for the DMN study: PrL (AP = 3.3 mm, ML = 0.8 mm, DV= 3.5 mm), Cg (AP = 1 mm, ML = 0.8 mm, DV = 2 mm), RSC (AP = −2.2 mm, ML = 0.7 mm, DV = 2 mm), and AI (AP = 3.2 mm, ML = 4.2 mm, DV = 3.5 mm) were used, and for the validation study: S1 (AP = 1 mm, ML = 4 mm, DV = 0.9 mm) was used. Next, four MR-compatible miniature brass screws (Item #94070A031, McMaster Carr, Atlanta, GA) were anchored to the skull, then multimode optical fibers (200 μM core; NA: 0.37) were chronically implanted to the experimental coordinates, and the surface of the skull was covered with dental cement to seal implanted components. Post operation analgesics included Bupivicane (2.5mg/ml s.c), Lidocaine (2.5% topical) and meloxicam (1.5mg/kg). Neomycin and polymyxin B sulfates and dexamethasone ophthalmic ointment, USP (Baushch & Lomb) was administered to prevent excessive dryness and infection of the eyes. All rats were allowed at least 1 week for recovery before any further experiment.

#### Fiber-photometry setup

A MRI-compatible, four-channel, spectrally resolved, fiber-photometry platform was used (Bao et al., 2020; Chao et al., 2022; Das et al., 2021; Meng et al., 2018; Oyarzabal et al., 2022; Zhang et al., 2022b, 2022a). We used interleaved 488 nm (OBIS Galaxy Laser 1236444, Coherent, Santa Clara, CA) and 405 nm (OBIS Galaxy Laser 1236439, Coherent, Santa Clara, CA) diode lasers for GCaMP signal excitation. The GCaMP signals derived from 488 and 405 nm excitation provide neuronal calcium activity and motion correction reference respectively. The interleaved lasers were launched into a dichroic mirror set (OBIS Galaxy Laser Beam Combiner, Coherent, Inc.), then equally split into 4 outputs by a 1-to-4 fan-out fiber optic bundle (BF42LS01, Thorlabs, Newton, NJ), and delivered into a fluorescence cube (DFM1, Thorlabs, Newton, NJ). Extra neutral density filters (NEK01, Thorlabs, Newton, NJ) were placed for additional adjustment of the final laser power before entering the fluorescence cube. The fluorescence cube contained a dichroic mirror (ZT405/488/561/640rpcv2, Chroma Technology Corp, Bellows Falls, VT) to reflect and launch the laser beam through an achromatic fiber port (PAFA-X-4-A, Thorlabs, Newton, NJ) into the core of a 105/125 mm core/cladding multi-mode optical fiber patch cable. The distal end of the patch cable was connected to an implantable optical fiber probe for both excitation laser delivery and emission fluorescence collection. The emission fluorescence collected from the fiber probe traveled back along the patch cable into the fluorescence cube, passed through the dichroic mirror and an emission filter (ZET405/488/561/640mv2, Chroma Technology Corp, Bellows Falls, VT), then was launched through an aspheric fiber port (PAF-SMA-11-A, Thorlabs, Newton, NJ) into the core of an AR-coated 200/230 mm core/cladding multi-mode patch cable (M200L02S-A, Thorlabs, Newton, NJ). The AR-coated multi-mode patch cable was connected to a spectrometer (QE Pro-FL, Ocean Optics, Largo, FL) for spectral data acquisition, which was operated by a UI software OceanView (Ocean Optics, Largo, FL). Laser outputs, oddball and control auditory stimuli, and spectrometer recordings were controlled using a programmable Med-PC® interface and synchronized with MRI using TTL pulses. The interleaved laser outputs were applied at a 50% duty cycle with 50 ms pulse duration. Emission spectra by the interleaved 488 and 405 nm lasers were collected at 1 ms after each laser switching with a 25 ms sampling window, resulting a 10 Hz effective sampling rate for both GCaMP signal time-course derived from 488 and 405 nm excitation. Functional MR images were acquired at 1 Hz, thus each fMRI timeframe covered 10 photometry data points.

#### Concurrent fMRI with fiber-photometry recordings

CBV-fMRI was acquired on a Bruker 9.4-Tesla/30-cm scanner with a BFG240-120 gradient insert. A homemade surface coil (1.2 cm inner diameter) with miniature circuit board served as an RF transceiver. Rats were orotracheally intubated and ventilated using a small animal MR-compatible ventilator (CWE Inc., MRI-1, Ardmore, PA). Under anesthesia by constant isoflurane (1.5–2%) blended with medical air, rats received tail vein catheters for intravenous drug and contrast agent injections, then were placed into a small animal cradle with a head-holder. Inside the cradle, a built-in circulating water line was linked to a temperature-adjustable water bath (Thermo Scientific, Waltham, MA) for stabilizing the rats’ body temperature, which was monitored using a rectal probe and maintained at 37 ± 0.5 °C. Mechanical ventilation volume and rate were adjusted to maintain EtCO2 of 2.8–3.2% and SpO2 above 90% as measured by a capnometer (Surgivet, Smith Medical, Waukesha, WI).

Before attaching the fiber-photometry patch cables to the implanted fiber ferrules on each rat, separate background spectra for the 405 and 408 nm lasers were acquired with the patch cable fiber tips pointing to a nonreflecting background in the dark MRI room. These background spectra were then subtracted during data analysis. Following setup processes, the cradle was pushed into MRI bore, and a bolus of dexmedetomidine (0.025 mg/kg; Dexdormitor, Orion, Espoo, Finland) cocktailed with paralytic agent rocuronium bromide (4.5 mg/kg; Sigma–Aldrich, St. Louis, MO) was injected via tail vein. Fifteen minutes after the bolus injection, continuous intravenous infusion of dexmedetomidine (0.05 mg/kg/h) and rocuronium bromide (9 mg/kg/h) cocktail was initiated and the isoflurane concentration was maintained at 0.5–1% for the remainder of the fMRI scanning session.

Magnetic field homogeneity was optimized first by global shim and followed by local first- and second-order shims according to the B0 map. Anatomical images for referencing were acquired using a rapid acquisition with relaxation enhancement (RARE) sequence (12 coronal slices, thickness = 1 mm, repetition time (TR) = 2500 ms, echo time (TE) = 33 ms, matrix size = 256 × 256, field-of-view (FOV) = 25.6 × 25.6 mm2, average = 8, RARE factor = 8). The slice center of the 5th slice from the anterior direction was aligned with the anterior commissure. Cerebral blood volume (CBV) fMRI scans were acquired using a multi-slice single-shot gradient echo echo-planar imaging sequence (GE-EPI) (slice thickness = 1 mm, TR = 1000ms, TE = 8.1 ms, matrix size = 80 × 80, FOV = 2.56 × 2.56 cm2, bandwidth = 250 kHz), with the same image slice geometry imported from the previously acquired T2-weighted anatomical image. For each session, a GE-EPI scan with 300 repetitions was performed prior to CBV fMRI scans, and rats were administered Feraheme (30 mg Fe/kg, i.v.) at approximately the 150th repetition. This data set provided the contrast before and after the Feraheme injection to verify that the injection was successful.

At the end of fMRI experiments, rats were administered atipamezole hydrochloride (3mg/kg, i.v.; ANTISEDAN, Orion, Espoo, Finland) for the reversal of the sedative and analgesic effects of dexmedetomidine, and sugammadex sodium (4 to 8mg/kg, i.v.; Merck Sharp & Dohme Corp., Kenilworth, New Jersey) for the reversal of the paralytic effect of rocuronium (Chao et al., 2018, 2022, Zhang et al., 2022b, 2022a).

#### Acoustic oddball task

Each acoustic oddball session was 20 min. During the session, control or oddball tones were pseudo-randomly given with an inter-stimulus interval of 2 seconds. We used 4 kHz and 8 kHz sinusoidal, 100 dB monophonic tones for control and oddball tone respectively, each tone was 50 ms long. Each session consisted of 580 controls and 20 oddballs, and a minimum initial sequence of 10 controls were given before pseudo-random presentation of oddballs, with a minimum of 10 control stimuli between any two oddballs.

#### Pre-processing of calcium fluorescence time series from fiber-photometry

To quantify GCaMP signal changes, the spectrum at each acquisition time point was fitted by the following function:

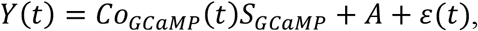

where *S_GCaMP_* represents the normalized reference emission spectra of GCaMP; *Co_GCaMP_*(*t*) is the unknown regression coefficients corresponding to the GCaMP signal; A is the unknown constant, and *ε*(*t*) is random error. The derived *Co_GCaMP_*(*t*) was detrended and mean-corrected. To measure the time-frequency energy distributions of GCaMP activity, *Co_GCaMP_*(*t*) was decomposed into a time-frequency function using the continuous wavelet transformation with complex Morlet wavelets as the mother wavelet. Based on the energy distributions of GCaMP activity, the GCaMP signals were band-pass filtered at 0.15-1.2 Hz for further analyses (**Figure 1C-F**).

#### Pre-processing of fMRI data

CBV-fMRI data was preprocessed with a standard pipeline using AFNI (Cox and Hyde, 1997). Specifically, for resting-state and oddball stimulation CBV-fMRI, preprocessing steps included despike, motion correction, skull stripping (Hsu et al., 2020), and spatial smoothing (full width at half-maximum, 0.6 mm). The CBV-fMRI images were normalized to a common 3D space aligned to a rat stereotaxic atlas (Lee et al., 2021). For resting-state CBV-fMRI data, the normalized images were linearly detrended and six-parameters of motion curves were regressed. Finally, independent component analysis was used to identify and remove physiological, movement, and thermal (machine) noise components (Griffanti et al., 2017; Rummel et al., 2013). All CBV-fMRI data (20 CBV-fMRI scans in 7 SD rats) were used to generate brain network components using a gICA (Smith et al., 2004). The number of components was set to 30 (Hsu et al., 2016; Jonckers et al., 2011), and the spatial distributions of individual components were identified using dual regression (Zuo et al., 2010). A one-sample t test was performed to generate the group component maps.

#### Histology

At the end of all experiments, rats were euthanized by a mixture of 1-2 ml of sodium pentobarbital and phenytoin sodium (Euthasol, Virbac AH, Inc., Westlake, TX), and transcardially perfused with saline followed by 10% formalin. The brains were removed and stored in 10% formalin overnight, then transferred into a 30% sucrose solution (in DI water) for 2–3 days, until brains sunk to bottom of storage bottles. These brains were cut into serial coronal sections (40 μm) using a freezing microtome (#HM450, Thermo Fisher Scientific, Waltham, MA) and mounted on glass slides. Fluoro-Gel II Mounting Medium (#17985-50, Electron Microscopy Sciences, Hatfield, PA) was covered on the brain slides to provide DAPI stain and for fluorescent imaging. Slides were imaged using a Zeiss LSM780 confocal microscope.

#### Bayesian switching dynamical systems (BSDS) model

To determine latent brain state dynamics underlying resting-state, we used a Bayesian Switching Dynamic Systems (BSDS) model (Taghia et al., 2018). Here we briefly describe the mathematical framework of the BSDS model. Let 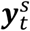 denote a D-dimensional vector of ROI timeseries obtained from subject *s* in time *t*, where *D* is the number of ROIs. Following the general formulation of the switching state-space models, we defined 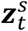 as the latent state variables and 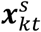 as the latent space variables associated to 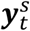 at the *k*-th latent state, that is 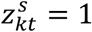. The 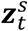 is a 1-of-*k* discrete vector with elements 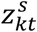, ∀*k* = 1,…,*K*. Two successive time instances are dependent through a 1st-order Markov chain of Hidden Markov Model (HMM). Using Markovian properties and given state transition probabilities ***A***, where 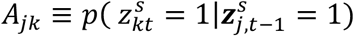 and a marginal distribution 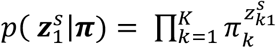 represented by a vector of initial probabilities ***π*** where 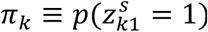, the probability distribution for the latent state variables is expressed by 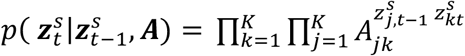 for all *t* > 1. We assume that at a given latent state *k* in time *t*, shown by 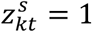, the observed vector 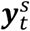 is generated via probabilistic interpretation of a factor analysis model (Everett, 2013; Ghahramani and Hinton, 1996) as:

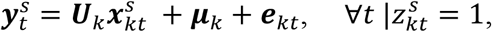

where *U_k_* is a *D* × *P* dimensional linear transformation matrix that transforms data to a subspace of lower dimensionality, *P* < *D*, described using a P-dimensional vector of latent space variables 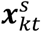 mediated by an overall bias ***μ**_k_* and a measurement noise ***e**_kt_*. With the normality assumption, that is 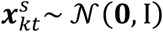 and 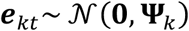, the marginal distribution of 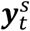 follows a Gaussian distribution as 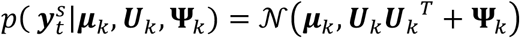 where T denotes the transpose operator. We then define a dynamical process on the latent space variables using an autoregressive (AR) model (Fox, 2009) of order *R* as:

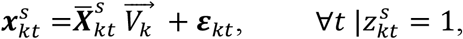

where 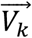 is a vector of AR coefficients. 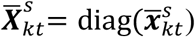 is a block diagonal isotropic matrix with elements of 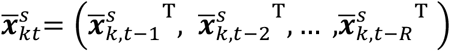 represented using latent space variables from the previous *R* time frames. 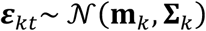 is the remaining error term in the latent space. All analyses conducted in this study use a 1st-order AR model (*R* = 1). Detailed theoretical derivations are provided in the previous study (Taghia et al., 2018).

We used a variational Bayesian (VB) framework to infer model parameters, including the number of brain states. The number of states is treated as a random variable, whose optimal value is learned from data using automatic relevance determination procedures implemented in a variational Bayesian framework. BSDS was initialized with 10 states, and it converged to 5 states.

BSDS estimated the posterior probability of each latent brain states at each time point and chose the latent brain state with the highest probability as the dominant state at that time point. Using the temporal evolution of the latent brain states, we measured temporal properties of each latent brain state, including occupancy rate and state switching probability. Occupancy rate quantifies the proportion of time that a state is chosen as the dominant state. State switching probability quantifies the chance that brain state at time point *t* either remains at its own state or switch to another brain state at the time point *t+1*. These temporal properties were examined to characterize their relationship with optogenetic stimulation conditions.

#### Functional connectivity of GCaMP signals and latent brain states

In this study, we used partial correlation as a measure of functional connectivity. Partial correlation estimates the correlation between any two brain regions after eliminating interdependencies on the common influences from other brain regions. Compared to Pearson correlation, partial correlation has been shown to more accurately reflect the relationships between brain regions (Marrelec et al., 2006). Using GCaMP signals of RSC, Cg, PrL, and AI, we computed the partial correlation between each pair of brain regions to estimate their time-averaged, static, functional connectivity. In BSDS analysis, each latent brain state is represented by a multivariate Gaussian distribution, which is described by the mean and covariance. To determine functional connectivity of each brain state, we computed the partial correlations from the estimated covariance matrices. For all the analyses, we conducted two-tailed *t*-test to check whether the correlation is significantly different from zero. Multiple comparison correction was implemented using false discovery rate (FDR) correction (*p* < 0.01).

#### Multivariate dynamical systems identification of causal interactions

MDSI is a state-space model consisting of a state equation to model the latent “neuronal–like” (quasi neuronal) states of the dynamic network (Ryali et al., 2011, 2016a, 2016b). MDSI estimates both intrinsic and experimentally modulated causal interactions between brain.

The state equation in MDSI is a multivariate linear difference equation or a first-order multivariate auto regressive (MVAR) model that defines the state dynamics as:

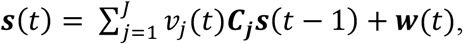

This equation represents the time evolution of latent neuronal signals in *M* brain regions, where *s*(*t*) is a *M* × 1 vector of latent signals at time *t* of *M* regions, ***C_j_*** is task-specific *M* × *M* connection matrix and *ν_j_*(*t*) is the *j*-th experimental condition at time *t*. ***C_j_***(*m,n*) denotes the strength of causal connection from *n*-th region to m-th region for the *j*-th task. ***w***(*t*) is a *M* × 1 state noise vector that is assumed to be Gaussian distribution with covariance matrix ***Q***(***w***(*t*)~*N*(0, *Q**I***)), where **I** is an identity matrix with size *M* × *M*. In addition, state noise vector at time instances 1, 2,…, *T*(***w***(1), ***w***(2),…, ***w***(*T*)) are assumed to be identical and independently distributed (iid). Estimating causal interactions between *M* regions specified in the model is equivalent to estimating the parameter ***C_j_***. In order to estimate ***C_j_*** the other unknown parameters, ***Q***, 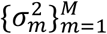, and the latent signal 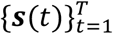 based on the observations 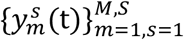, *t* = 1, 2,.. *T*, where *T* is the total number of time samples and *S* is number of subjects, needs to be estimated.

In resting-state analysis, we estimated causal interaction between AI, Cg, PrL, and RSC during resting-state by letting *ν_j_*(*t*) = 1 for all time points, where the experimental condition, *j*, equals to 1. In oddball experiment, we estimated causal interaction between brain regions associated with oddball (*j* = 1) and control (*j* = 2) stimulus trials by modeling *ν*_1_(*t*) equals to 1 for 4 seconds after each oddball stimulus trial and equals to 0 in rest of the time points, and modeling *ν*_2_(*t*) as the opposite of *ν*_1_(*t*). MDSI estimated strength of dynamic causal interaction per connection per condition. A paired *t*-test was used to examine whether the strength of dynamic causal interaction between conditions is different and multiple comparison correction was implemented using false discovery rate (FDR) correction (*p* < 0.01)

## Notes

### Competing Interest Statement

The authors have declared no competing interest.

