## Supplementary Information for "Neuronal dynamics of the default mode network and anterior insular cortex: Intrinsic properties and modulation by salient stimuli"

### Supplementary Figures

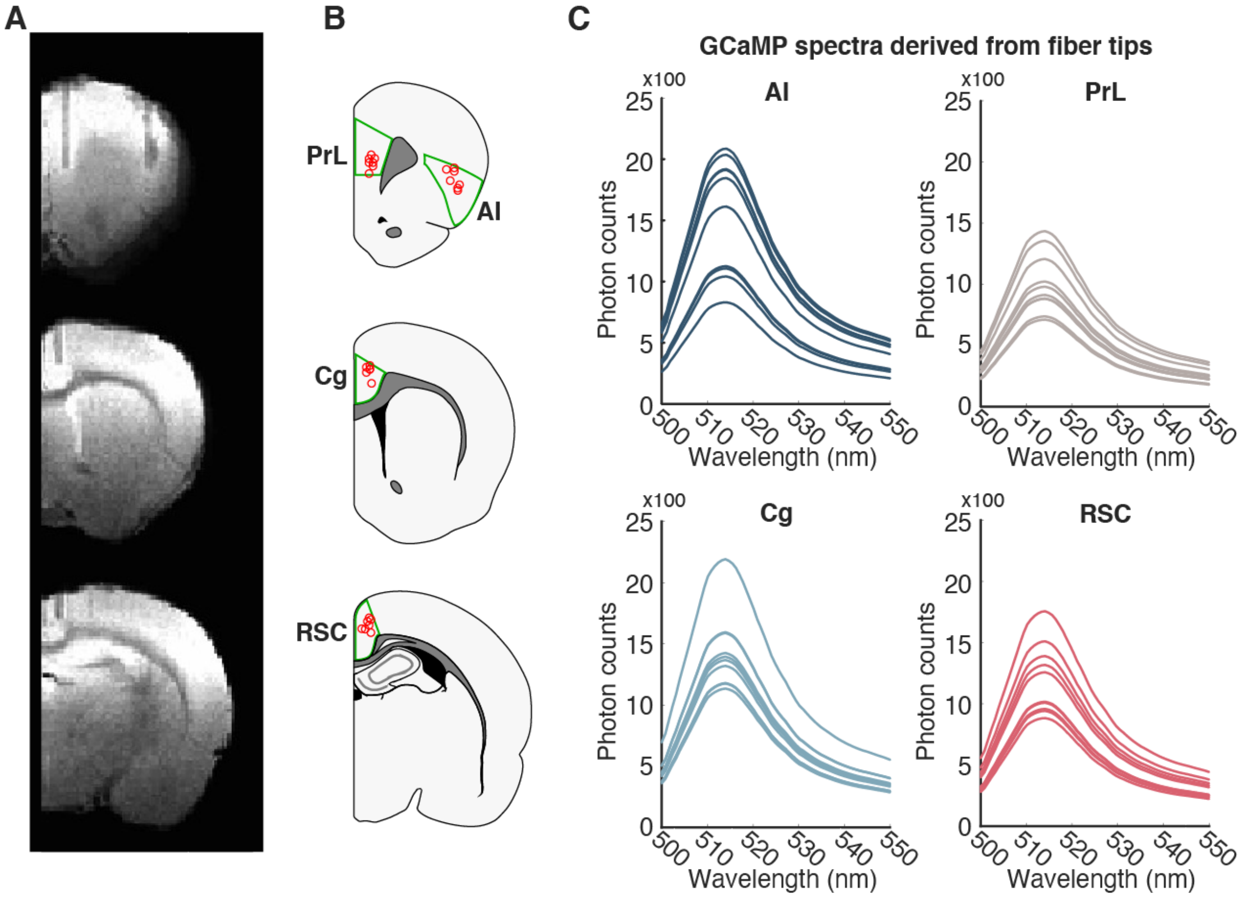


**Figure S1. Validation of fiber implant locations and GCaMP signals (A)** Fiber implant locations were identified from the anatomical T2 MR images of each rat. **(B)** Summary of fiber tip locations in all individual rats are labeled in red circles, anatomical binderies for the brain regions of interest are outlined in green. **(C)** GCaMP expression at the fiber tips were validated with fiber-photometry detection of GCaMP spectrum.

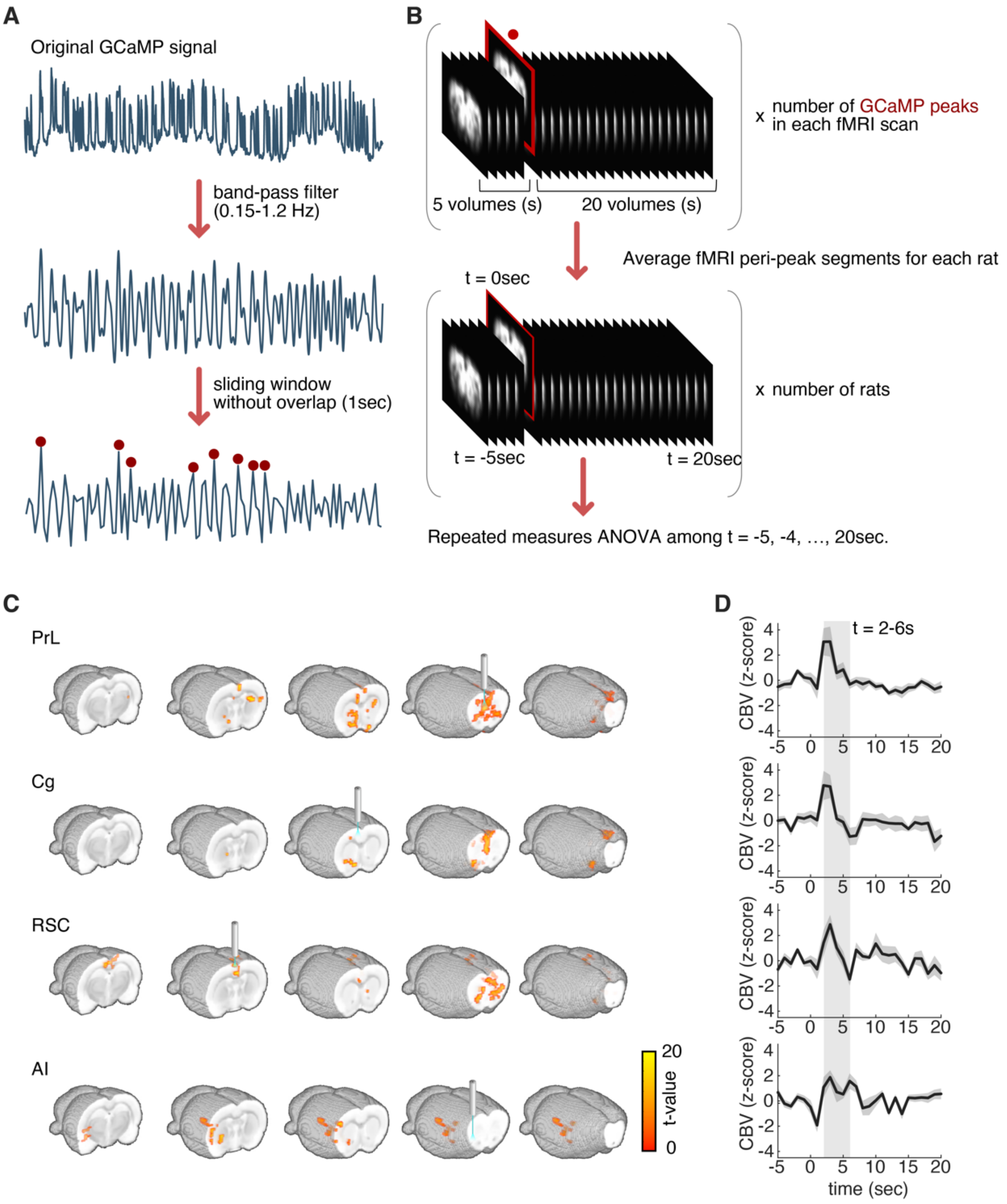


**Figure S2. Peri-event CBV-fMRI responses peaked at ~3 s after GCaMP spikes. (A)** To derive CBV-fMRI spatial and temporal response patterns from local GCaMP signals, we first band- passed the local GCaMP signals with cutoff frequencies at 0.15 Hz and 1.2 Hz, then we summed the data points across every 10 acquisitions (at 10 Hz) to match the temporal resolution of CBV-fMRI (1 Hz). **(B)** We extracted the timings of neural activation peaks from the processed GCaMP time courses, and then cropped the corresponding CBV-fMRI time series data from -5 s to +20 s of those peak timing. Next, we averaged the cropped CBV-fMRI time series intra-individually, then performed group-level repeated measures ANOVA among the CBV-fMRI time. **(C)** Statistical maps of CBV-fMRI responses to neuronal activation in PrL, Cg, RSC and AI ($p < 0.05$, n = 16). **(D)** Averaged dynamic CBV change time courses extracted from the global CBV activation maps in **(C)** (n = 16).

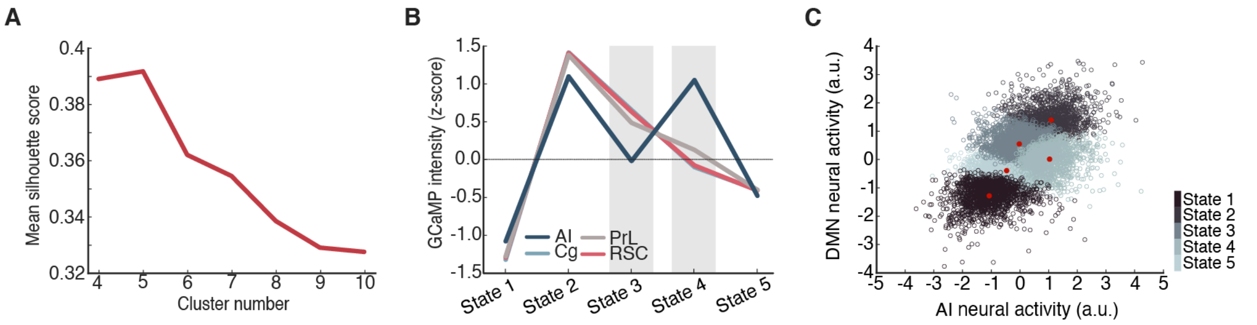


**Figure S3. Distinct brain states derived from GCaMP activity patterns across DMN-related nodes reveal network topology in global fMRI signals. (A)** The silhouette value when k = 4 to 10 in k-means clustering. k = 5 yields the maximum silhouette value. **(B)** Normalized GCaMP intensity across states identified from K-means analysis. **(C)** Since the activities of all putative DMN regions were comparable across states, we consolidated the 4 dimensions (AI, Cg, PrL, and RSC) into 2 dimensions by using the average activity of the DMN nodes as one dimension and using the AI activity as another dimension, and visualized the cluster distribution of all 5 states with a 2-D scatterplot. The red dots indicate the centroid of each state.

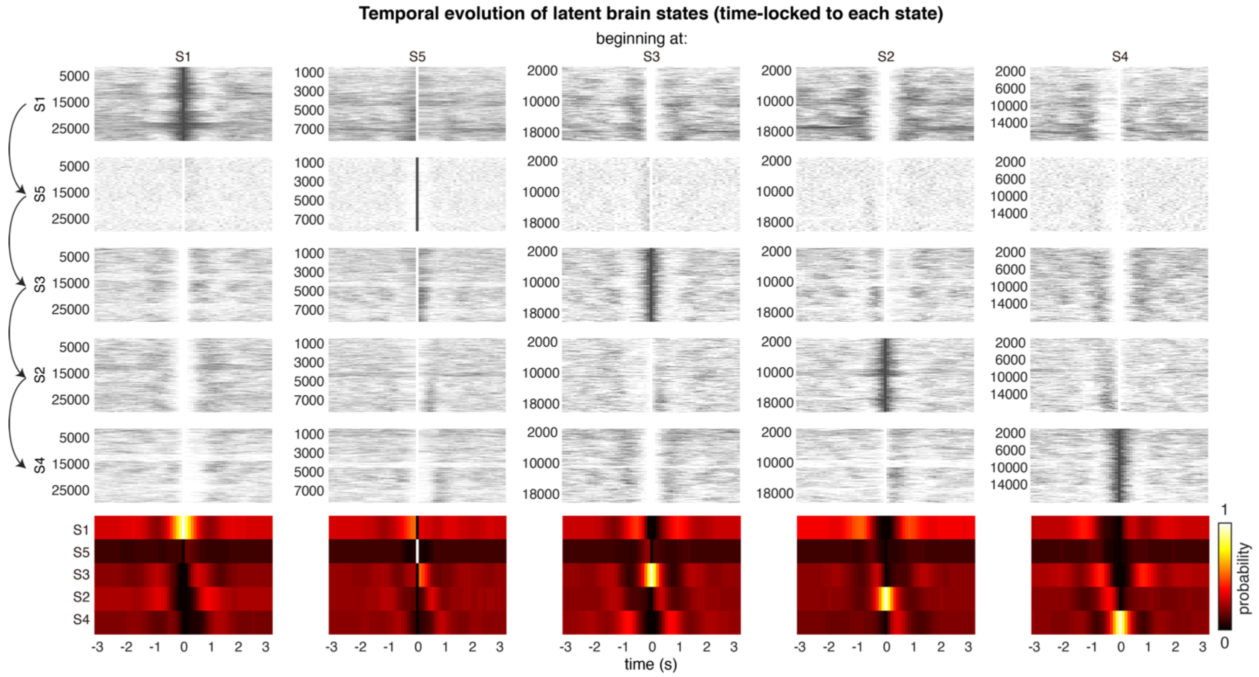


**Figure S4. Temporal evolution of latent brain states revealed that state transition follows a cyclic pattern.** Each column represents the temporal evolution of latent brain states that time-locked to a specific latent brain state. The bottom colormap summarizes the occupancy probability of latent brain states in respect to each time-locked latent brain state.
